# A scalable approach for tissue microarray profiling using spatial single cell transcriptomics

**DOI:** 10.1101/2025.08.08.669400

**Authors:** Claudia Arnedo-Pac, Jo Heffer, Marina Golotiuk, Ania M. Piskorz, Sarah J. Aitken

## Abstract

Spatially resolved transcriptomics was named Method of the Year 2020^1^ and has continued to evolve rapidly since then, providing novel insights in development, physiology, and disease processes. Many approaches now offer excellent performance in formalin-fixed paraffin embedded tissue, opening up a wealth of archived clinical samples to these high-throughput spatial biology modalities. In this Chapter, we introduce a methodology to apply 10x Xenium *In Situ* to FFPE tissue microarrays. Using this method we have achieved a cost-effective approach to profile hundreds of genes with subcellular resolution in over a hundred human liver samples. Our method can easily be adapted to other tumour types, diseases, or model systems.

## Introduction

Spatial transcriptomics methods can be broadly classified into (i) imaging-based and (ii) sequencing-based approaches^2^. Imaging-based methods directly visualise gene expression within tissue sections, often at single-cell or subcellular resolution, and may be low throughput (e.g. such as fluorescence *in situ* hybridisation (FISH)) or higher throughput (e.g. Xenium (10x Genomics), MERFISH (Vizgen), and COSmx (Bruker)), but can be limited by the multiplexing of targeted gene probes. In contrast, sequencing-based methods (e.g. Visium SD and HD (10x Genomics) and GeoMx Digital Spatial Profiler (NanoString)) capture and sequence RNA from tissue sections, providing higher throughput and whole transcriptome coverage, but accurate cell segmentation may be challenging and some approaches lack single cell resolution. The choice of method depends on the required resolution, throughput, number of genes, and tissue type. In this Chapter, we describe our method to apply Xenium *In Situ* from 10x Genomics (10x) to achieve transcriptome analysis at subcellular resolution over a large slide area for application to tissue microarrays (TMAs) and compatibility with formalin-fixed paraffin-embedded (FFPE) liver tissue from clinical samples (**Fig. 1-3**).

**Figure 1.**
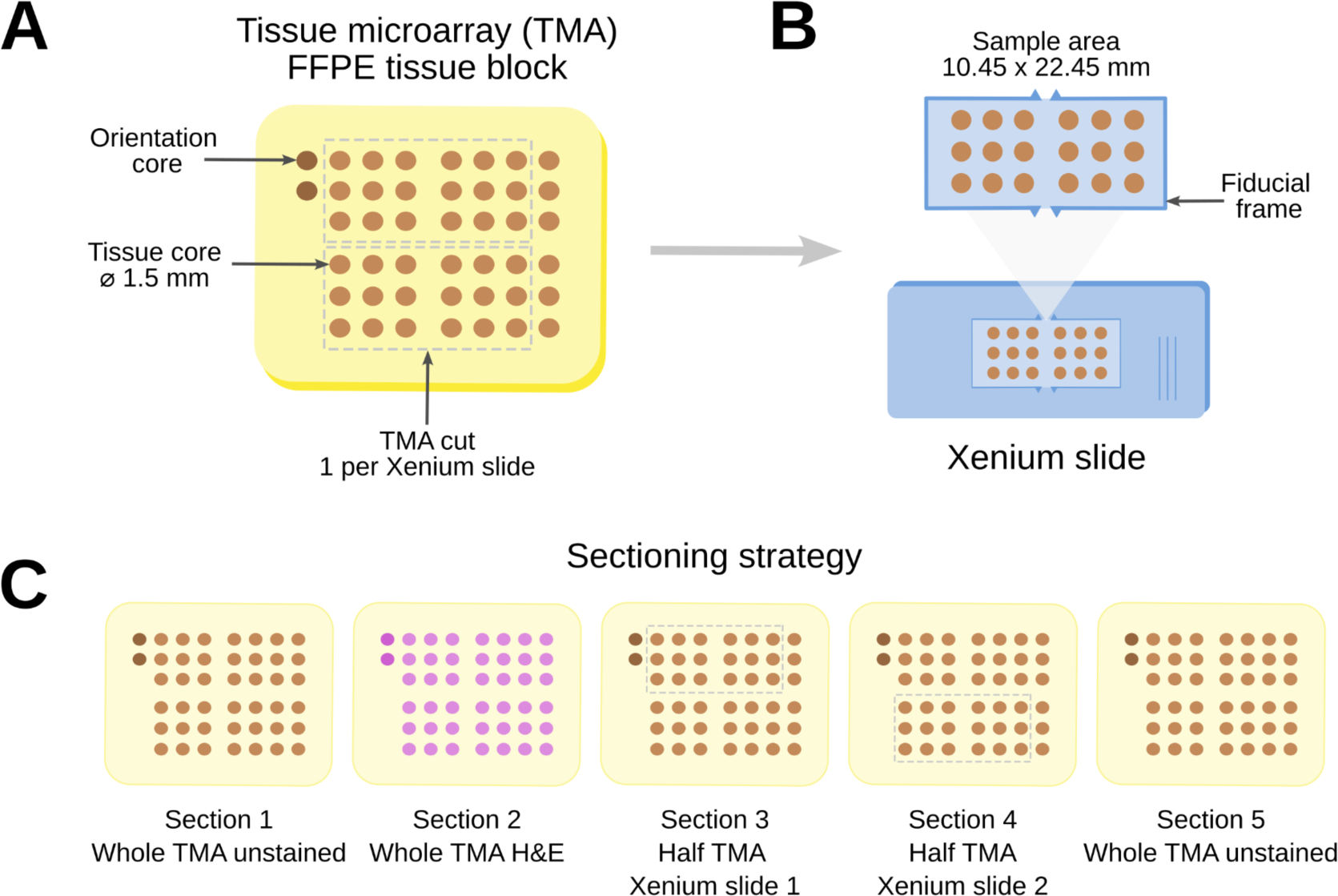
Preparation of tissue microarrays for Xenium *In Situ*. **(A)** Design of our TMA blocks containing 42 liver tissue cores and 2 orientation cores at the top left. Given the dimensions of the sample area in Xenium slides (shown in panel B), the TMAs were divided in two and the last column of liver cores was excluded for analysis. **B)** Example of the TMA sample area within the Xenium slide. Note that the sample area does not include the fiducial frame surrounding it. To avoid inaccuracies within Xenium workflow, it is strongly recommended to avoid tissue touching the fiducial frame. **C)** Schematic of the sectioning strategy for our TMAs. For each TMA, we generated two sections for Xenium (one TMA section per Xenium slide, highlighted by the grey dashed line), one adjacent section for an H&E reference, and unstained sections at the front and back for optional downstream validation experiments (e.g. immunohistochemistry or immunofluorescence).

The Xenium *In Situ* workflow allows quantitative, high-resolution (single cell) *in situ* detection of hundreds of genes in parallel (**Fig. 2**)^3^. After de-paraffinisation and de-crosslinking of FFPE tissue (or fixation and permeabilisation if using FF tissue), the panel of probes is added to the Xenium slide. Each probe has two RNA hybridisation targets, allowing circularisation by ligation, with a region that encodes a gene-specific barcode. The circular template undergoes enzymatic rolling circle amplification, which generates multiple copies of the gene-specific barcode for each RNA target. There is an option to add the Multimodal Cell Segmentation workflow^4^. This approach allows accurate cell segmentation by combining nuclear (DAPI staining), membrane (ATP1A1, E-cadherin, CD45 antibodies), cytoplasmic (18S ribosomal RNA), and cytoskeleton (alphaSMA, vimentin antibodies) markers with the 10x nuclear expansion algorithms. Following autofluorescence quenching, the Xenium slide is loaded onto the Xenium Analyzer instrument, which performs multiple cycles of fluorescent probe hybridisation, imaging, and probe removal^5^. This creates optical signatures specific for each gene, allowing the identification, location, and quantification of each transcript target. The data are immediately available for quality control (QC) and visualisation as part of the Xenium Onboard Analysis pipeline and Xenium Explorer software (**Fig. 3**).

**Figure 2.**
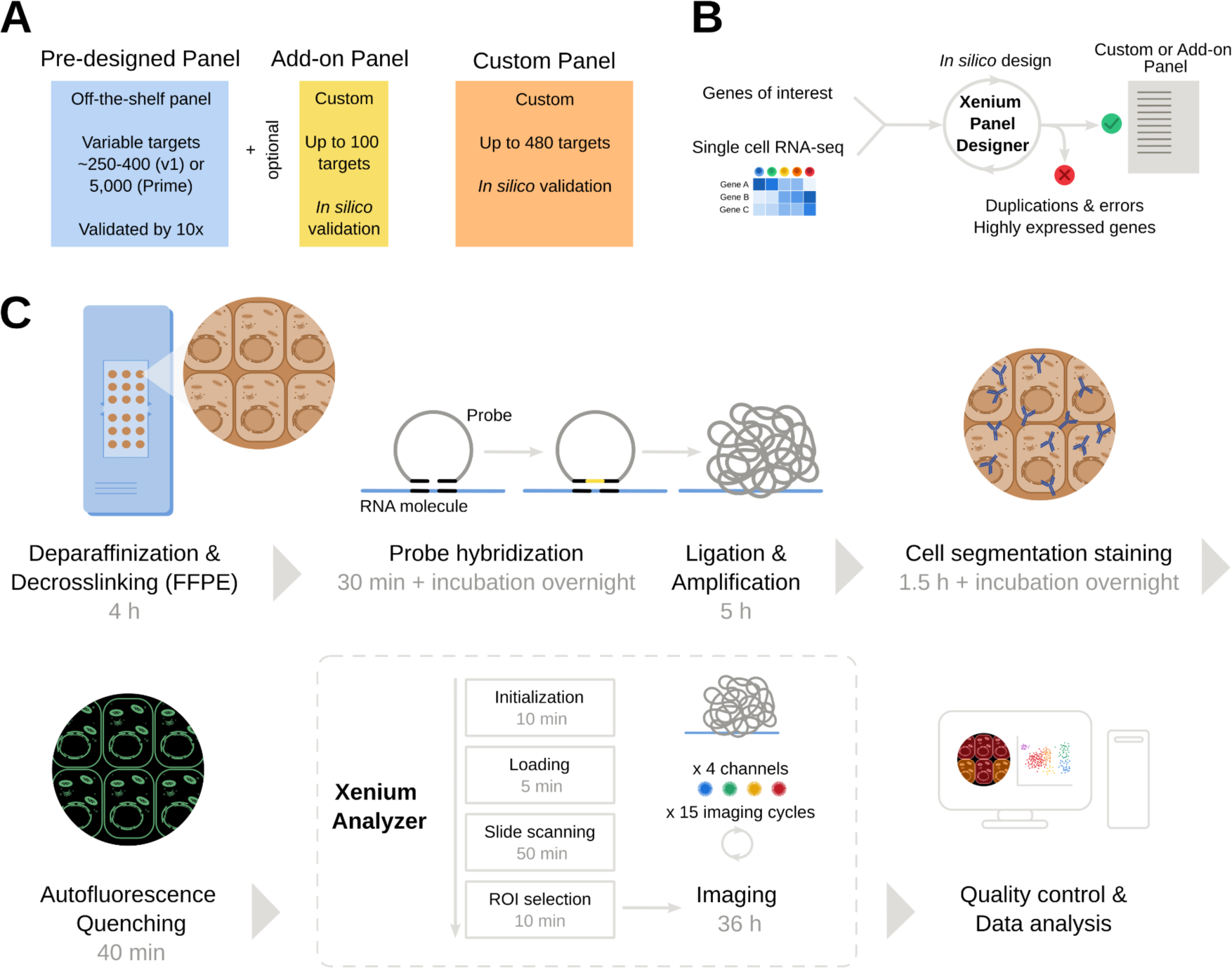
Xenium *In Situ* panel design and workflow. **(A)** Xenium panel design options. **(B)** Schematic of the custom or add-on panel design process. Starting with a list of genes of interest and tissue-matched single cell RNA-sequencing data, the Xenium Panel Designer application optimises the panel design and flags errors and highly expressed genes to be excluded from the panel. **(C)** Schematic of Xenium *In Situ* workflow for FFPE tissue. Note, timings are approximate.

To increase Xenium throughput and cost-effectiveness, here we have analysed FFPE TMAs containing multiple liver samples from over one hundred donors. TMAs are paraffin blocks which are constructed to contain dozens to hundreds of FFPE tissue cores from separate histopathology paraffin blocks, allowing tissue from multiple patients to be sampled on one slide (**Fig. 1A**). This multiplexing approach allows time- and cost-efficient analysis of clinical samples and limits batch effects. Since many clinical researchers and cancer biologists have already constructed TMAs with patient cohorts of interest, there are now exciting new opportunities (and challenges) to open up these archived resources to high-throughput spatial biology approaches.

In this Chapter, we describe our approach to tissue sectioning (to fit the bespoke Xenium slide), gene panel design, and offer tips for troubleshooting the Xenium protocol and Multimodal Cell Segmentation workflow, including QC and downstream analysis (see ‘Notes’ section). The Xenium materials and methods described below are the most up-to-date version of 10x Genomics protocols^3–6^ at the time of writing; these are updated periodically, and we recommend that readers check for the most recent version.

## Materials

### Histology materials

All equipment that will come into contact with the paraffin section should be cleaned thoroughly with RNaseZap. The inside of the water bath should also be cleaned before filling.

1. TMA tissue block

- The size and composition of tissue microarrays (TMAs) will vary between users. For the project detailed in this chapter, the TMA tissue blocks measured 21 x 15 mm. Each TMA contained 42 FFPE tissue cores measuring 1.5 mm diameter with inter-core distance of 1.5 mm. Two orientation cores were located at the left top corner of the block (**Fig. 1A**).
2. TMA sectioning

- Rotary microtome
- Water bath filled with Ultrapure water (approximately 45°C)
- Ice tray
- Forceps
3. Xenium slide and adjacent normal slide mounting and pre-Xenium storage

- Xenium slides (room temperature). Since Xenium slides are stored at −20°C, they need to be removed from the freezer to equilibrate to room temperature for at least 30 min prior to sectioning.
- Razor blade
- Cutting mat
- Lamp
- Xenium template slide. This is created by tracing the sample area on a normal glass slide, using the template provided in the Tissue Preparation Guide^6^. Note that the available area to place the sectioned tissue in a Xenium slide, referred to as “sample area”, measures 10.45 x 22.45 mm (**Fig. 1B**).
- Superfrost Plus slides
- Slide rack
- Oven set at 42°C
- Desiccator
4. Xenium slide preparation and deparaffinisation

- Oven set at 60°C
- Xylene: 2 baths
- 100% ethanol: 2 baths
- 96% ethanol: 2 baths
- 70% ethanol
- Nuclease free water
5. Haematoxylin and eosin (H&E) staining for post-Xenium slide and adjacent normal slide

- Quencher removal solution: add 17.4 mg Sodium Hydrosulfite to 10 ml Milli-Q water and vortex for 10 seconds
- Autostainer (e.g. Leica ST5020) programmed with a standard FFPE H&E protocol and coverslipper (e.g. Leica CV5030)
6. H&E scanning for post-Xenium slide and adjacent normal slide

- Brightfield slide scanner (e.g. Leica Aperio AT2)
- Digital pathology slide management software (e.g. Leica Aperio eSlide Manager software)
7. Post-Xenium slide and adjacent normal slide storage

- Desiccator

### Xenium workflow materials

The reagents and equipment required for the Xenium workflow, including probe hybridisation, cell segmentation staining, and imaging are presented below. Two slides are prepared in parallel during each run, hence the volumes below stated are for two reactions (i.e. two slides).

1. Xenium slides (x2) and sample preparation reagents:

- Perm enzyme B
- Decrosslinking buffer
- Xenium probe hybridization buffer
- Xenium post-hybridization wash buffer
- Xenium probe hybridization buffer
- Xenium ligation buffer
- Xenium ligation enzyme A
- Xenium ligation enzyme B
- Xenium amplification mix
- Xenium amplification enzyme
2. Xenium cassette kit (2 cassettes, 16 lids)
3. Optional: Xenium pre-designed gene expression probes

- In this project, we used the Xenium human Multi-Tissue and Cancer Panel
4. Optional: Xenium custom probes

- In this project, we used a 51 to 100 custom add-on gene panel (see “Xenium panel design” below)
5. Optional: Xenium cell segmentation staining reagents:

- Xenium block and stain buffer
- Xenium multi tissue stain mix
- Xenium stain enhancer
- Xenium cassette insert and slide seals
6. Xenium decoding reagents (1 run, 2 slides):

- Xenium decoding reagent module A
- Xenium decoding reagent module B
7. Xenium cell segmentation detection reagents (module)
8. Xenium decoding consumables (1 run, 2 slides)
9. Buffers and solutions:

- Ethanol 100%
- Ethanol 70%
- Nuclease-free water
- PBS 10X (phosphate buffered saline) pH 7.4, RNase-free. Freshly prepare PBS 1X for each run, using nuclease-free water)
- 10% Tween-20
- PBS-T: 1X (phosphate buffered saline pH 7.4, 10% Tween-20. Prepare fresh for each run)
- Urea solution, 8 M
- TE buffer1X (Tris-EDTA) pH 8.0
- DMSO (dimethyl sulfoxide), molecular biology grade 100%
- KCl (potassium chloride), RNase-free
- 70% isopropanol
10. Equipment:

- Xenium Analyzer (software v.3.2.1.2)
- Xenium Thermocycler Adaptor
- Thermal cycler (e.g. BioRad C1000 Touch with 96-deep well reaction module).
- Thermomixer
- Vortex
- Mini centrifuge
- Refrigerated microcentrifuge
- Centrifuge with plate rotor
- Forceps
- Lint-free laboratory wipes
- Graduated cylinders, glass bottles with cap (sterile)
11. Master mixes:

a. Decrosslinking and probe hybridisation.

- Diluted perm enzyme B 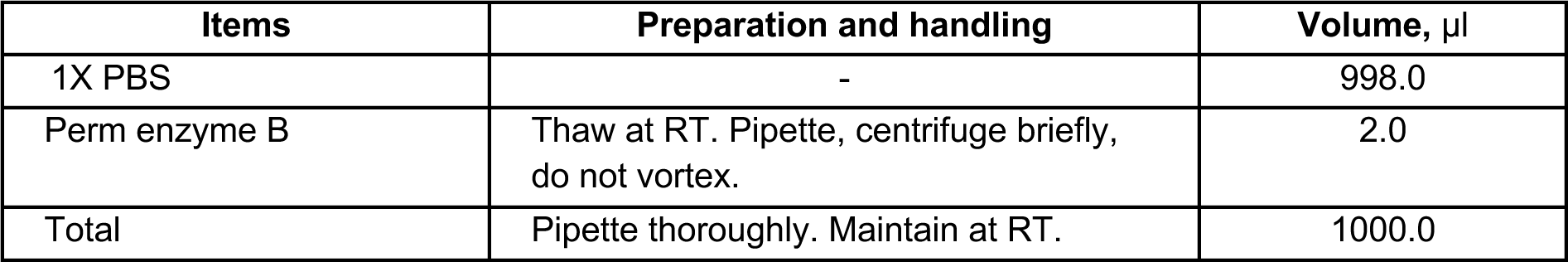
- Decrosslinking buffer 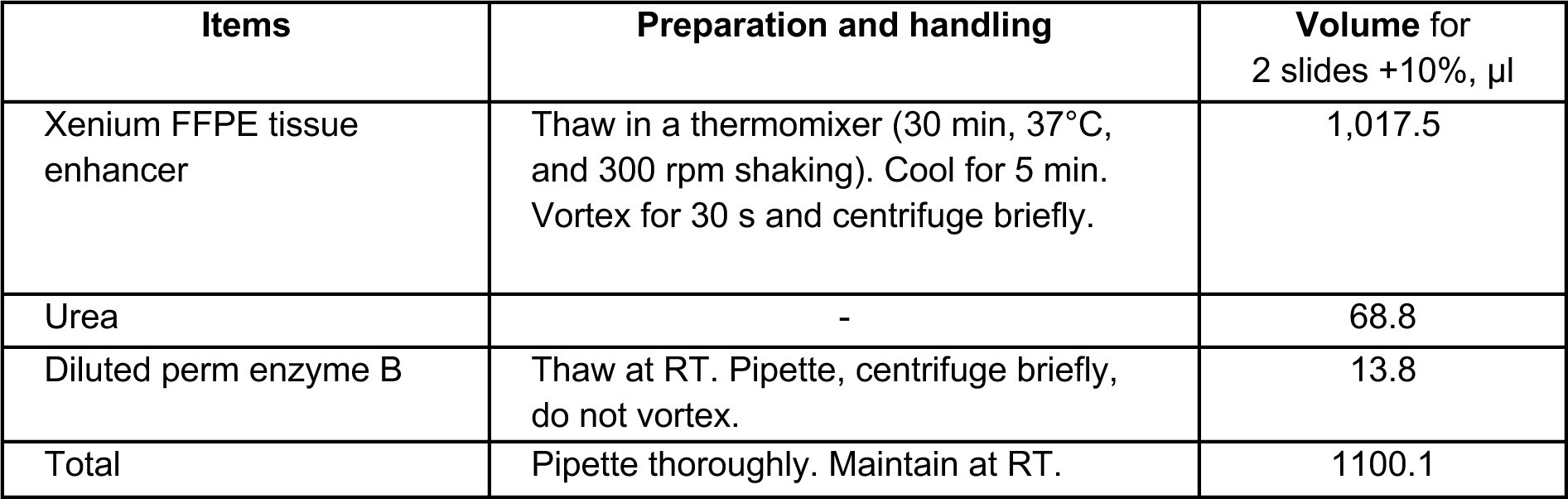
- Probe hybridization mix (add-on custom probe panels used with pre-designed probe panels) 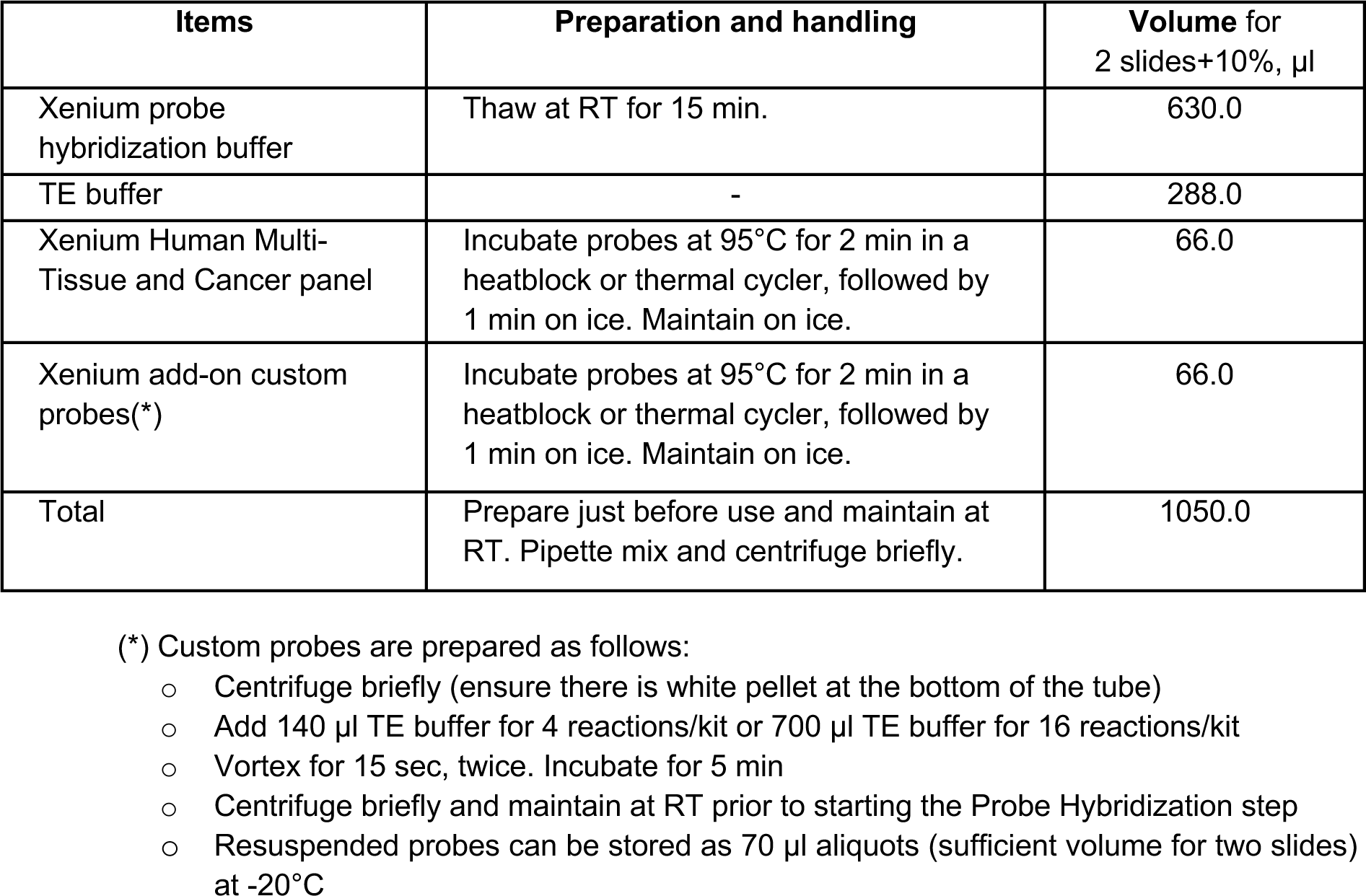
b. Ligation

- Ligation mix 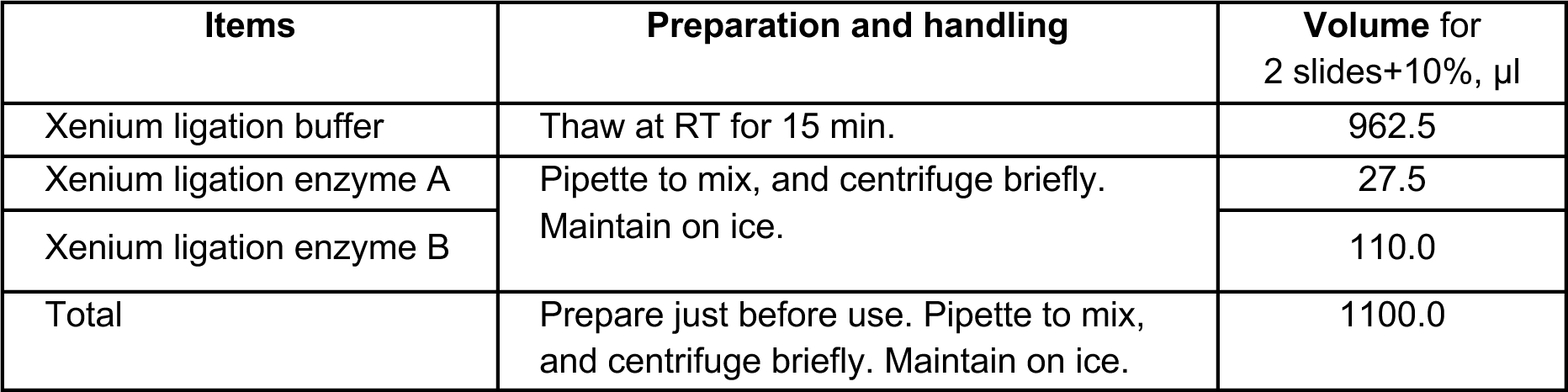
c. Amplification

- Amplification master mix 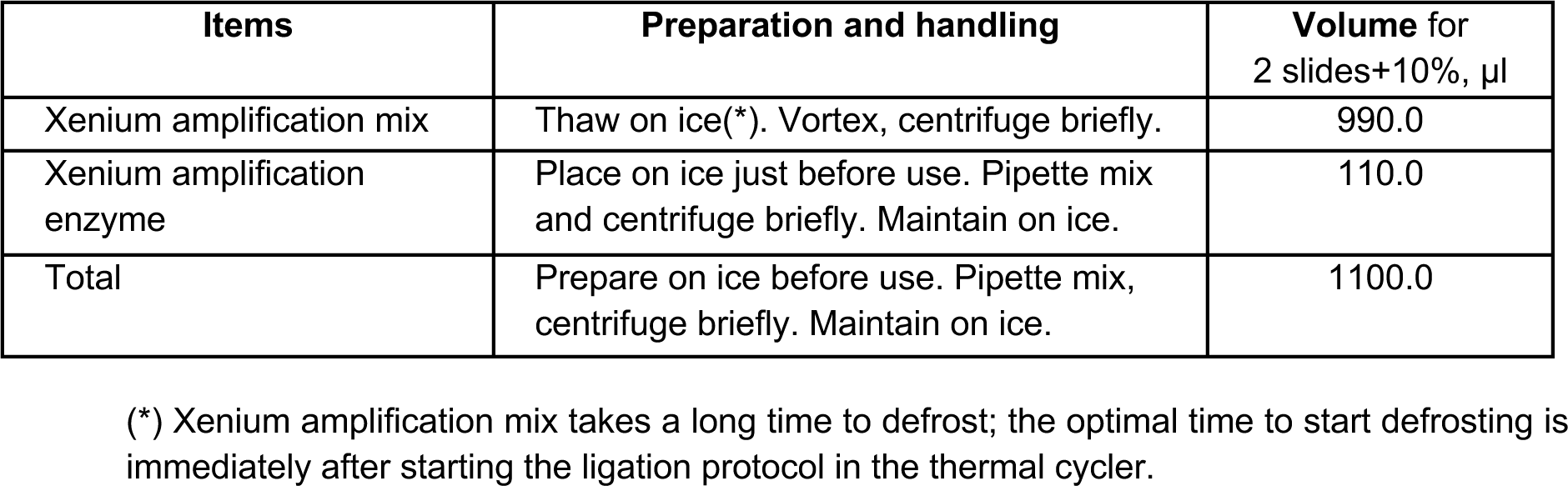
d. Optional: Cell segmentation staining

- Diluted xenium block and stain buffer 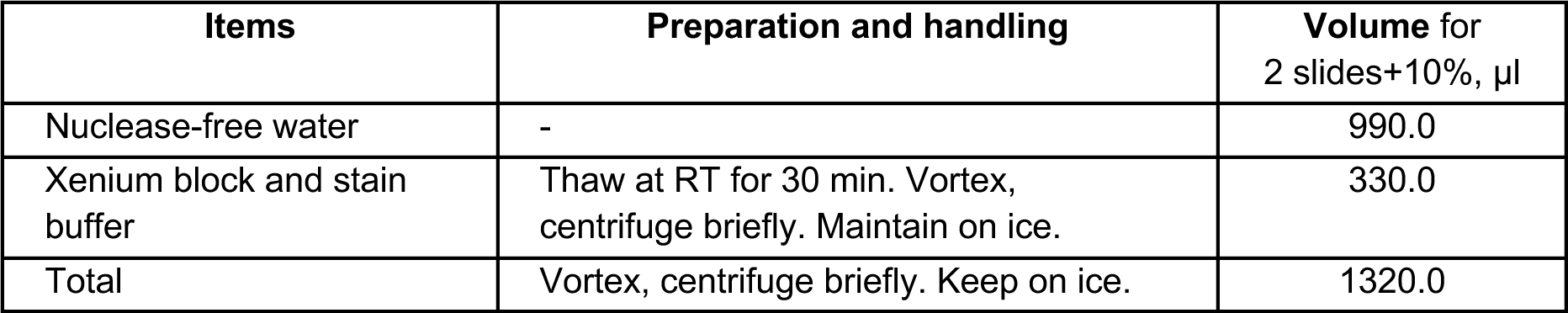
e. Autofluorescence quenching

- Diluted reducing agent B 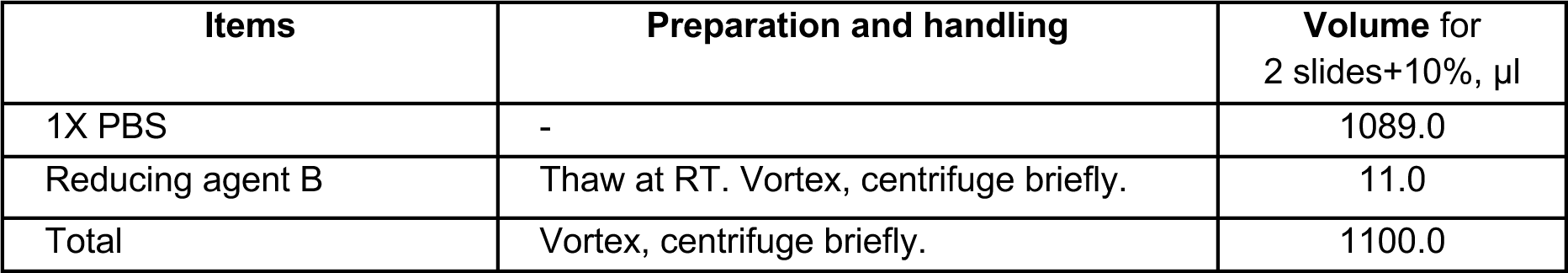
- Autofluorescence Solution 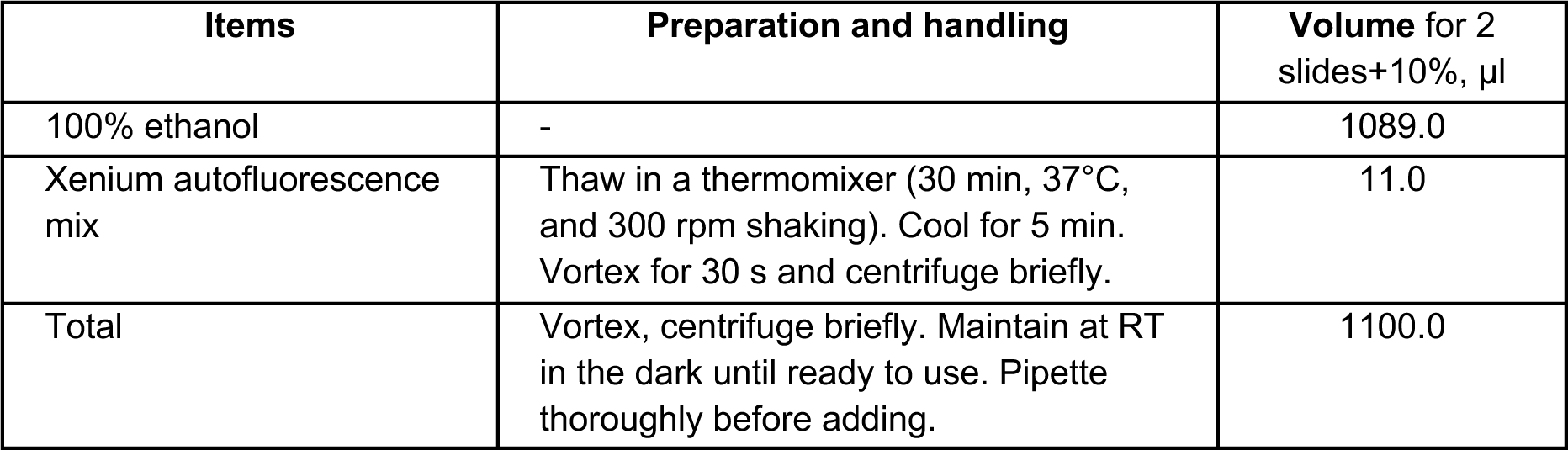
f. Xenium Imaging

- Xenium Instrument Wash Buffer (bottle #1) 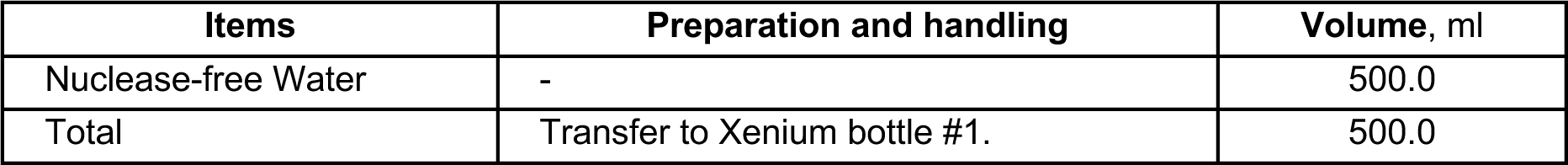
- Xenium Sample Wash Buffer A (bottle #2) 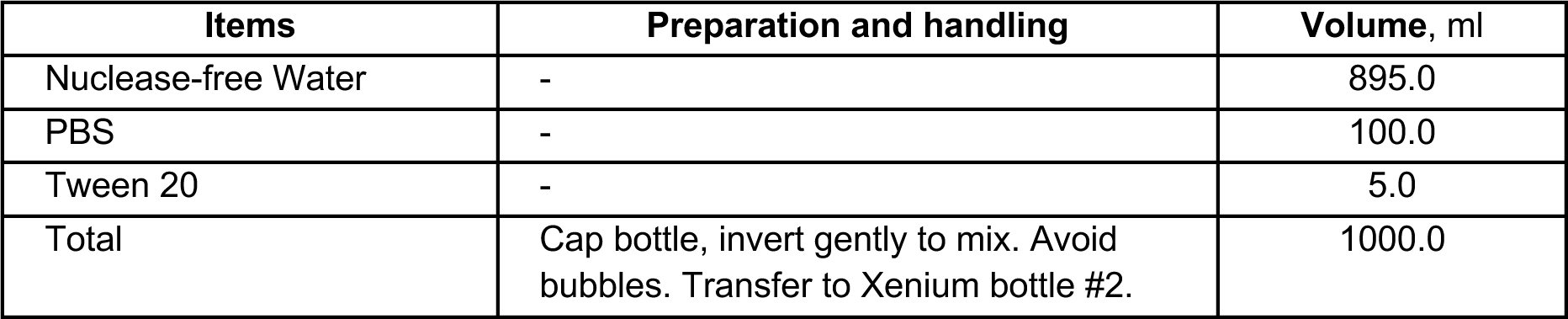
- Xenium Sample Wash Buffer B (bottle #3) 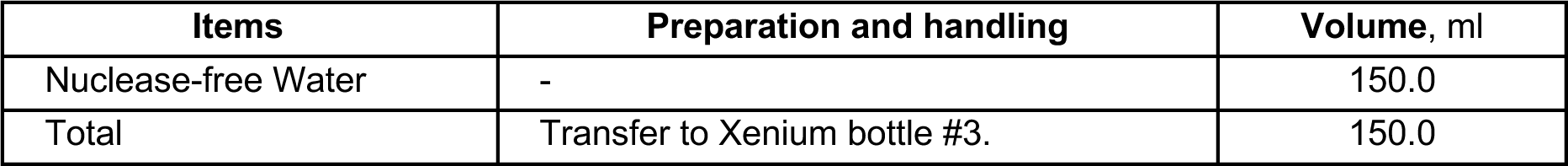
- Xenium Probe Removal Buffer (bottle #4) 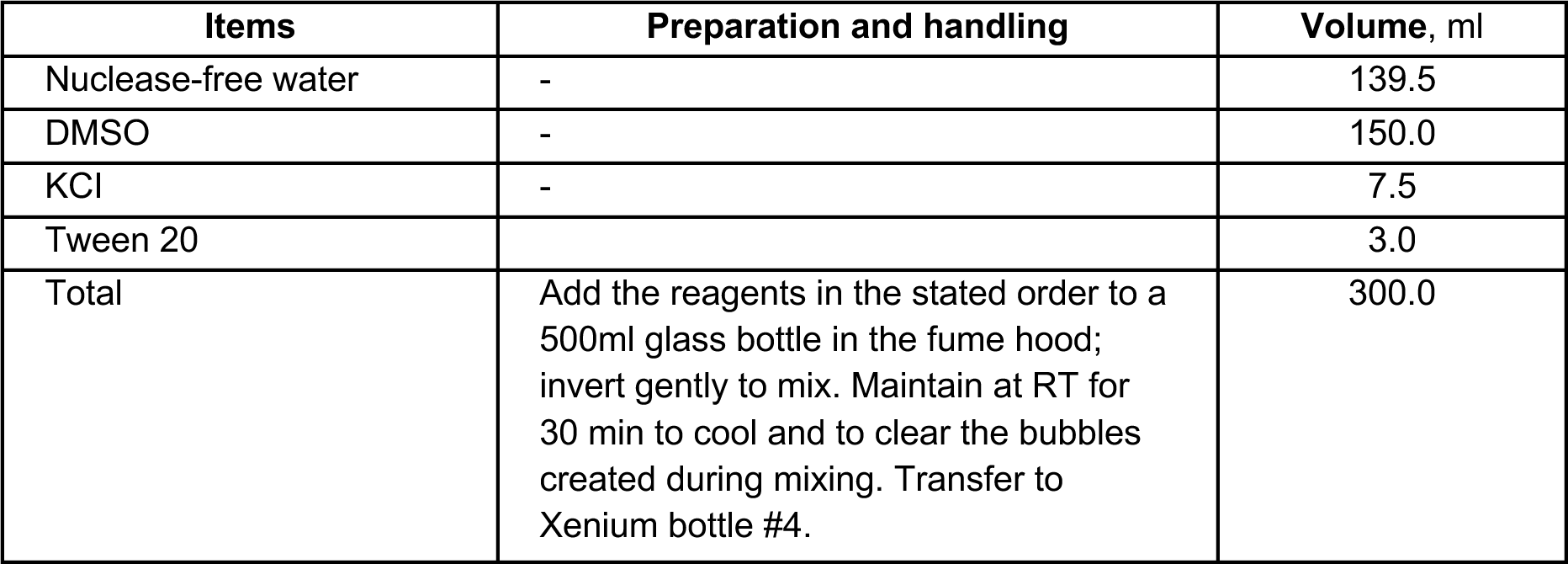

## Methods

### 1. Xenium panel design

*In situ* RNA expression detected by 10x Xenium is based on the identification of fluorescent signals from a panel of gene probes targeting multiple transcripts of interest. Currently, 10x offers 3 main types of gene panels: (1) pre-designed gene panels, (2) custom gene panels, and (3) add-on custom panels to complement pre-designed panels. Pre-designed panels have been validated by 10x, whereas custom and add-on gene panels designed by the user through the 10x Xenium Panel Designer web application do not have experimental validation provided by 10x. In order to support decision making of this crucial step, the three main types of Xenium gene panels are briefly described here (**Fig. 2A**).

1. **Pre-designed gene panels**. These off-the-shelf panels have been developed and tested by 10x. Pre-designed panels comprise Xenium v1 and Xenium Prime 5K panels. Xenium v1 panels, containing between 247 and 380 target genes, cover the main cell types of diverse human or mouse tissues. Xenium Prime 5K panels are more comprehensive, targeting 5,000 genes in the human or mouse transcriptome. Pre-designed panels can be explored at <u>10xgenomics.com/products/xenium-panels</u>. Panel files, including gene names, transcript IDs, target cell types, probe sequences in FASTA format and genomic coordinates in BED format, are freely available to download. These panels have a short delivery time because they do not need bespoke manufacturing.
2. **Custom gene panels**. These panels are fully customisable by the user, allowing up to 480 target genes at the time of writing. The panel design is carried out through Xenium Panel Designer web application running on the 10x Genomics Cloud (no installation required). Xenium Panel Designer helps to design the optimal distribution of probes across target genes of interest based on reference single cell expression data and provides an *in silico* prediction of the panel performance, as described below. Following panel design, custom panels are made to order by 10x.
3. **Add-on custom panels**. These smaller custom panels are designed to work in combination with a pre-designed gene panel (also referred to as base panel). They can be used to expand the basic panel by up to 100 genes. They can also be used for other purposes such as targeting different gene isoforms or nucleotide variants within mRNA molecules (also referred to as advanced add-on panels). As with custom gene panels, they are designed through Xenium Panel Designer and made to order. Note that add-on custom panels are designed to be used in combination with the user-specified base panel (i.e., they are not recommended to be used with a different base panel).

In this experiment, we used the pre-designed Xenium Human Multi-Tissue and Cancer Gene Expression Panel (Xenium v1) together with a 100 gene add-on panel. In order to design the add-on panel, we followed the standard workflow within 10x Xenium Panel Designer (**Fig. 2B**):

1. **Gene list drafting.** Our add-on panel aimed to incorporate 100 additional genes to the Human Multi-Tissue and Cancer base panel. In order to prioritise which genes would be included in the add-on panel, we first selected a broad range of genes relevant for our specific experiment (e.g. liver expressed genes). We then checked the presence of these genes in the pre-designed base panel Xenium Human Multi-Tissue and Cancer Gene Expression Panel. Those included in the base panel were removed from our draft list of add-on genes. We curated and prioritised this gene list until we obtained 110 genes with potential interest to be included in the add-on panel. This analysis was carried out independently of the Xenium Panel Designer platform.
2. **Utilising the 10x Xenium Panel Designer.** We logged in to the Xenium Panel Designer application at <u>cloud.10xgenomics.com/xenium-panel-designer</u> and started a new panel design request. The application allows saving the design in progress, thus the work can be completed through multiple sessions.
3. **Panel information**. The application first asks for basic metadata, including: material type (formalin-fixed paraffin-embedded (FFPE) or fresh frozen (FF)), organism (human or mouse), reference genome (GRCh38, mm10, or non-reference genome), the specific base panel among the different 10x pre-designed gene panels, and the size of the add-on panel (either 1-50 or 51-100 genes).
4. **Reference datasets**. Users need to specify up to 5 single cell RNA-sequencing (scRNA-seq) reference datasets matching the tissue type of interest. This is a crucial step because these datasets are used by the 10x algorithms to model the expected gene expression of the target genes across cell types to generate the optimal add-on panel design. Discrepancies between the reference datasets used in panel design and the actual tissue where the panel is applied could cause reduced Xenium workflow performance. Reference scRNA-seq data can be obtained from different sources: 1) a variety of publicly available datasets pre-built within Xenium Panel Designer; 2) in-house datasets uploaded by the user; 3) a combination of 1 and 2. In our case, we chose publicly available liver scRNA-seq datasets accessible within Xenium Panel Designer^7^.
5. **Designing the add-on panel**. From this point onwards, the design becomes an iterative process, where the user modifies a list of genes they wish to include in the add-on panel and runs Xenium Panel Designer to make automated *in silico* predictions about the performance of the proposed gene panel. This process contains the following parts:

a. *Submission of the gene list*. Xenium Panel Designer requests to introduce a gene list containing HUGO Gene Nomenclature Committee (HGNC) canonical gene names or Ensembl IDs into the application. Optionally, the number of probes per gene can also be specified in this step. 10x suggests starting the panel design with 10 or 20 genes in addition to the maximum number of gene slots available in the add-on panel (i.e. 110-120 genes for the 100 gene panel). The genes need to be ranked from high to low priority, as Xenium Panel Designer will introduce them to the add-on panel following the priority list up until the maximum number of gene slots available in the add-on panel (N) is reached. Genes beyond N will be excluded unless any of the top-N genes failed Xenium Panel Designer workflow checks, in which case subsequent genes beyond N will be used.
b. *Review of gene names and IDs*. When the gene list is submitted, the algorithm first validates that the gene names are correctly mapped to the reference genome data. Incorrect or duplicated gene symbols or IDs are flagged in this step. If needed, users can go back to the previous step and modify the gene list accordingly.
c. *Panel design and recommendations*. This step contains the most relevant information to assess the add-on panel design. The Panel Design Summary contains:

- Warnings. A summary of warning flags followed by proposed solutions will appear at the top.
- Performance of the panel across cell types in the reference datasets (Panel Utilization per Cell Type). Xenium workflow is based on the *in situ* identification of fluorescent signals across different gene probes. Of note, the number of fluorescent signals that can be resolved per tissue area or volume is finite. Transcripts from highly expressed genes (per cell or across cells; e.g., marker or housekeeping genes) can yield closely located fluorescent signals that are difficult to deconvolute to individual transcripts (phenomenon referred to as optical crowding). It is therefore important to consider the detection budget in the tissue, and ensure that the combined predicted fluorescent signals from the base and add-on panels do not exceed this theoretical threshold. The Panel Utilization per Cell Type will summarise this information among the cell types annotated in the reference datasets. The ideal scenario would be a panel where the expression for each cell type is within the detection budget, thus decreasing the probability of optical crowding and increasing the sensitivity for transcript detection.
- Panel Utilization for Highly Expressed Genes. Similar to the previous plot, the Panel Utilization for Highly Expressed Genes will show whether the detection budget per gene is within the optimal range across cell types in the reference datasets. This can be used to flag highly expressed genes in a given cell type or ubiquitously expressed genes across different cell types, which could cause optical crowding. The use of highly expressed genes should be avoided because of the risk of optical crowding. For example, in our case, the first run of the process identified an hepatocyte marker, albumin (*ALB*), as a highly expressed gene in hepatocytes. Given that our panel contained other hepatocyte marker genes, we decided to remove *ALB* from the panel to avoid potential optical crowding. An alternative strategy is to retain highly expressed genes in the panel, but to decrease the number of probes targeting them (see below).
- Probeset Summary. This is a visual representation of the number of probes (probeset) generated for each gene. By default, the algorithm allocates 8 probes per gene; the higher the number of probes, the higher the sensitivity. As mentioned above, while it is recommended to exclude highly expressed genes from the panel, an alternative strategy is to retain highly expressed genes by decreasing the number of probes targeting them. However, given that the performance of add-on custom panels will not be experimentally validated during the process of panel design, the risk of optical crowding cannot be excluded.
- Expression Heatmap. This plot shows the normalised scRNA-seq expression of add-on genes across cell types in the reference datasets. It is useful to predict how the add-on panel will perform in the tissue of interest (e.g., by checking if known marker genes are expressed in the expected cell types).
- Excluded genes. Genes in the base and/or add-on panels that are not included in the reference scRNA-seq datasets will raise a warning and will be discarded from the add-on panel design, as their reference expression cannot be analysed. Based on this summary, and the recommendations provided by Xenium Panel Designer, the user can iterate the process by providing a new gene list to verify until there are no issues with the panel design.
d. *Panel submission*. Once the panel has been finalised and submitted, the user can download the genes and associated Ensembl IDs in the panel, a BED file containing probe coordinates, a JSON file with full panel data and metadata required by Xenium Analyzer to run the workflow (see Section 5), and an HTML summary of the panel design and recommendations.

### 2. TMA Sectioning

The number of serial sections required will vary according to the experimental design (**Note 1**); here we use one section for H&E, two sections for Xenium (half TMA section per Xenium slide), and additional sections for validation assays (**Fig. 1C**). In our design, the adjacent H&E section allowed us to track the orientation of the TMA after the TMA was cut to fit the Xenium sample area. Pre-Xenium sectioning optimisation may be required (**Note 2**).

1. Prepare for sectioning by putting the block on ice, filling the water bath and heating to 45°C, removing the Xenium slide from the freezer, and turning 2 ovens on, one at 42°C and one at 60°C.
2. Use a microtome to take serial sections at 5 μm. Ensure sections are perfectly consecutive serials to help with mapping and annotation of tissue cores between the Xenium slide and adjacent whole TMA section.
3. Lay the serial sections on a cutting mat and assess for quality, pay specific attention to any rolled up or missing cores. If quality is low take another set of serial sections and check again until the required quality is achieved (**Note 3**).

### 3. Xenium slide and adjacent normal slide mounting

For simplicity, we will refer here only to the tissue sections used for H&E staining (adjacent normal section; section 1) and Xenium workflow (sections 2 and 3) (**Fig. 1C**).

1. Use the razor blade to separate the serial set into individual sections; it can help to chill the razor blade on the ice tray to prevent it sticking to the sections.
2. Using forceps, transfer section 1 to the water bath, leaving it to float on the water until it is flat and free of wrinkles (**Note 4**). This is the *adjacent normal* section.
3. Collect the *adjacent normal* section onto a Superfrost slide and leave to air dry.
4. Use the razor blade to cut out the required region from the **second** section, the top half of the TMA, to fit the Xenium sample area (**Note 5**) (**Fig. 1C**).
5. Optional step: hold the Xenium template slide above the section to check it will fit entirely within the fiducial frame (**Fig. 1B**); this is not necessary if the cut region is much smaller than the frame. Note that the fiducials need to be visible to Xenium Analyzer in order to carry out a Xenium run. Overlapping tissue or any other material that avoids the correct detection of the fiducials could generate downstream issues.
6. Using forceps transfer the top half section to the water bath, allow it to float, and collect it onto the Xenium slide within the fiducial frame, maintaining TMA tissue orientation at all times. In case the fiducials are covered, 10x suggests re-sectioning the tissue and using a new Xenium slide.
7. Repeat steps 4-6, this time using the **third** serial section to collect the bottom half of the TMA (**Fig. 1C**).

### 4. Slide drying and storage before Xenium workflow

1. Air dry all slides for 30 min at room temperature.
2. Bake the *adjacent normal* slide for 1 hour at 60°C then proceed to H&E staining. If necessary, the slide can be stored at room temperature prior to staining.
3. Bake the Xenium slides for 3 hours at 42°C and then dry overnight in a desiccator. Proceed to the Xenium workflow. Note that slides can be stored for up to 4 weeks before carrying out the Xenium workflow. However, it is recommended to minimise the time between this step and running Xenium, as the RNA quality may decrease over time, thus decreasing Xenium workflow sensitivity.

### 5. Xenium workflow

Xenium slide preparation involves several steps (**Note 6**): deparaffinisation and decrosslinking of FFPE tissue (or fixation and permeabilisation if using FF tissue), probe hybridisation, ligation, amplification, cell segmentation staining (optional step for accurate segmentation of cell membranes, nuclei, and cytoplasm), and autofluorescence quenching (**Note 7**). Following this, the Xenium Analyzer performs multiple cycles of fluorescent probe hybridisation, imaging, and probe removal^5^. This creates optical signatures specific for each probeset (gene), allowing the identification, location, and quantification of each transcript target (**Fig. 2C**).

#### 3.1 Deparaffinisation, decrosslinking and probe hybridisation

1. Bake Xenium slides for 2 hours at 60°C and then start the deparaffinisation process by incubation with xylene (twice), followed by rehydration in graded ethanols, 100%, 96% and 70%; finally place the slides in nuclease free water.
2. After deparaffinisation, place the Xenium slides with TMA sections into the Xenium cassettes. Using clean forceps, remove the slide from the jar with nuclease free water. Use a lint-free laboratory wipe to remove any remaining water. Place the slide in the bottom of the cassette with the tissue facing up. Apply the top of the cassette. Ensure the slide is correctly positioned at the bottom of the cassette. Otherwise, when you apply the top part, the slide may crack. Finally, check there are no gaps between the top and bottom of the cassette.
3. Add 500 µl 1X PBS and immediately proceed to step 4.
4. Prepare Decrosslinking buffer. Xenium FFPE Tissue Enhancer should be thawed in a thermomixer for 30 min (it is advisable to start preparing this before proceeding to the decrosslinking step).
5. Remove 1X PBS from step 3. Add 500 µl decrosslinking buffer and apply the new Xenium cassette lids to the cassettes.
6. Place the cassettes on the thermocycler adaptor on the thermal cycler. Start the incubation protocol (Table 1) (**Note 8**).
7. Once the decrosslinking protocol is complete, place the cassettes on a clean work surface, remove and discard the lids. Proceed immediately to the next step.
8. Remove the decrosslinking buffer.
9. Wash step: add 500 µl PBS-T to the well and incubate for 1 min. Remove all PBS-T.
10. Repeat the previous wash step two more times (three in total).
11. Add 500 µl PBS-T to the well and proceed to the next step.
12. Prepare Probe Hybridization Mix.
13. Aspirate all PBS-T from step 11 and carefully add 500 µl Probe Hybridization Mix (important not to introduce bubbles). Apply a new Xenium cassette lid.
14. Place the cassettes on the Thermocycler Adaptor on the thermal cycler, start the probe hybridization protocol protocol (Table 2).
15. Remove the Xenium cassettes from the thermal cycler (21 hour incubation), remove and discard the lids.
16. Remove the probe hybridisation mix from the slide by aspirating from the corner of the cassette; ensure you do not touch the tissue. It is important to work quickly so that the tissue does not cool down after hybridization.
17. Wash step: immediately add 500 µl PBS-T and incubate 1 min. Remove all PBS-T and repeat the wash one more time. The accuracy of the post-hybridisation washes is critical to achieving good quality data.
18. Prepare the thermal cycler for the next protocol by starting the ‘pre-equilibration’ step (Table 3).
19. Add 500 μl Xenium post hybridisation wash buffer. Apply a new Xenium cassette lid.
20. Place the cassettes on the Thermocycler adaptor on the thermal cycler, and proceed to the ‘post-hybridisation’ step of the incubation protocol (Table 3).
21. Immediately proceed to the next step.

**Table 1.** Decrosslinking protocol (for thermal cycler C1000 Touch, Bio-Rad)

| Lid temperature | Reaction volume |  |
| --- | --- | --- |
| 80°C | 100µL |  |
| Step | Temperature | Run time (hh:mm:ss) |
| Hold | 22°C | Hold |
| Decrosslinking | 80°C | 00:30:00 |
| Re-equilibrate | 22°C | 00:10:00 |
| Hold | 22°C | Hold |

**Table 2.** Probe hybridization protocol (for thermal cycler C1000 Touch, Bio-Rad)

| Lid temperature | Reaction volume |  |
| --- | --- | --- |
| 50°C | 100µL |  |
| Step | Temperature | Run time (hh:mm:ss) |
| Pre-equilibrate | 50°C | Hold |
| Probe Hybridization | 50°C | Overnight (16-24h) |
| Hold | 50°C | Hold |

**Table 3.** Post hybridization wash protocol (for thermal cycler C1000 Touch, Bio-Rad)

| Lid temperature | Reaction volume |  |
| --- | --- | --- |
| 37°C | 100µL |  |
| Step | Temperature | Run time (hh:mm:ss) |
| Pre-equilibrate | 37°C | Hold |
| Post hybridisation | 37°C | 00:30:00 |
| Hold | 37°C | Hold |

#### 3.2 Ligation

1. Prepare the ligation mix in advance. Pipette the mix and centrifuge briefly. Maintain on ice.
2. Remove the Xenium cassettes from the thermal cycler, remove and discard the cassette lids.
3. Remove all of the buffer from the well corners.
4. Immediately add 500 µl PBS-T and incubate 1 min. Remove all PBS-T and repeat this step two more times (three in total).
5. Add 500 µl ligation mix. Apply a new Xenium cassette lid.
6. Place the cassettes on the Thermocycler adaptor on the thermal cycler, start the ligation protocol (Table 4).
7. After the Ligation protocol is complete, immediately proceed to the next step.

**Table 4.** Ligation protocol (for thermal cycler C1000 Touch, Bio-Rad)

| Lid temperature | Reaction volume |  |
| --- | --- | --- |
| 37°C | 100µL |  |
| Step | Temperature | Run time (hh:mm:ss) |
| Pre-equilibrate | 37°C | Hold |
| Ligation | 37°C | 02:00:00 |
| Hold | 37°C | Hold |

#### 3.3 Amplification

1. Prepare amplification master mix.
2. Remove the Xenium cassettes from the thermal cycler, remove and discard cassette lids.
3. Remove all of the buffer from the well corners.
4. Wash step: immediately add 500 µl PBS-T and incubate 1 min. Remove all PBS-T and repeat this step two more times (three in total). The accuracy of post-ligation washes is critical for performance.
5. Add 500 µl amplification master mix. Apply a new Xenium cassette lid.
6. Place the cassettes on the Thermocycler adaptor on the thermal cycler, start the amplification protocol (Table 5).
7. Remove the Xenium cassettes from the thermal cycler, remove and discard the cassette lids.
8. Remove Amplification master mix.
9. Immediately add 500 µl TE buffer and incubate for 1 min. Remove all TE the buffer and repeat this wash step two more times (three in total). Do not remove the last 500 µl TE buffer. Apply a new Xenium cassette lid.
10. Prepare the staining buffer preparation while keeping the slides in cassettes at RT.

**Table 5.** Amplification protocol (for thermal cycler C1000 Touch, Bio-Rad)

| Lid temperature | Reaction volume |  |
| --- | --- | --- |
| 30°C | 100µL |  |
| Step | Temperature | Run time (hh:mm:ss) |
| Pre-equilibrate | 30°C | Hold |
| Amplification | 30°C | 02:00:00 |

#### 3.4 Optional step: Multimodal cell Segmentation Staining

1. Remove the cassette lids (do not discard, keep for the next steps). Remove all TE buffer.
2. Wash with 1000 µl 70% ethanol and incubate at 2 min RT. Remove the ethanol.
3. Wash with 1000 µl 100% ethanol and incubate at 2 min RT. Remove the ethanol. Repeat one more time.
4. Wash with 1000 µl 70% ethanol and incubate at 2 min RT. Remove the ethanol.
5. Add 1000 µl PBS-T quickly to avoid overdrying the tissue.
6. Remove all PBS-T and add 500 µl diluted Xenium block and stain buffer. Do not discard the remaining volume of the buffer.
7. Apply the Xenium lids from step 1. Incubate for 1 hour at RT.
8. During the incubation step, add 220 µl diluted Xenium block and stain buffer (remaining from step 6) to the Xenium multi-tissue stain mix tube. Pipette mix, centrifuge briefly. Incubate for 30 min at RT. Centrifuge for 10 min at 4°C (14,000 rcf). Maintain on ice.
9. Remove the cassette lids (do not discard). Remove the buffer from step 6. Using forceps place the Xenium cassette insert onto the Xenium cassette. Take extra care during the cell segmentation step when applying the insert. To avoid damaging the tissue, ensure that the insert does not come into contact with the tissue. To avoid over drying the tissue do not remove the buffer from the second slide before placing the insert on the first slide.
10. Add 100 µl from Xenium multi-tissue stain mix tube (avoid the pellet) to the Xenium Cassette using the cut-out to pipette solution under the Xenium cassette insert.Bubbles may affect performance of the assay. To avoid this, do not press the pipette to the second stop, leaving 2-3 µl of buffer inside the tip. Pipette slowly. If bubbles are observed, gently remove the insert, pierce the bubbles (with a 100 µl tip) without touching the tissue, and carefully replace the insert.
11. Apply the slide seal and press it along the edges of the cassette.
12. Incubate the cassettes overnight (16-24 h) at 4°C, in the dark.
13. Before the incubation will end, thaw Xenium stain enhancer at RT for 10 min. Centrifuge for 5 s (ensure white particulate is at the bottom of the tube). Add 1,100 μl 1X PBS, pipette mix and centrifuge again.
14. Remove slide seal from cassettes. Slowly add 200 μl PBS-T into the insert cut-out to float the inserts. Using forceps, carefully remove inserts and discard.
15. Remove all mix.
16. Immediately add 500 µl PBS-T and incubate for 1 min. Remove all PBS-T and repeat this step two more times (three in total).
17. Add 500 μl Xenium stain enhancer (from step 13).
18. Apply the cassette lids from step 9. Incubate for 20 min at RT.
19. Remove the cassette lids. Remove all stain enhancer from step 17.
20. Immediately add 500 µl PBS-T and incubate for 1 min. Remove all PBS-T and repeat this step one more time (two in total).
21. Add 500 μl PBS-T and proceed immediately to the next step.

#### 3.5 Autofluorescence Quenching and Nuclei Staining

1. Remove all PBS-T from the well.
2. Add 500 μl diluted reducing agent B, apply the new cassette lids and incubate for 10 min at RT.
3. Remove the cassette lids (do not discard). Remove mix from the well from step 2.
4. Wash with 1,000 μl 70% ethanol. Incubate for 1 min at RT. Remove the ethanol.
5. Wash with 1,000 μl 100% ethanol. Incubate for 1 min at RT. Remove the ethanol. Repeat this step one more time (two in total).
6. Add 500 μl autofluorescence solution to the well.
7. Reapply the cassette lids and incubate for 10 min at RT in the dark.
8. Remove and discard the cassette lids. Remove all of the autofluorescence solution.
9. Add 1,000 μl of 100% ethanol. Incubate for 2 min at RT. Remove the ethanol. Repeat two more times (three in total).
10. Place the cassettes (without lids) to dry on the Thermocycler adaptor on the thermal cycler, start the drying protocol (Table 6). The thermal cycler lid must stay open.
11. After drying immediately remove the Xenium cassettes and add 1,000 μl 1X PBS. Incubate for 1 min at RT in the dark.
12. Remove all 1X PBS from the well.
13. Add 1,000 μl PBS-T and incubate for 2 min at RT in the dark.
14. Remove all PBS-T.
15. Add 500 μl Xenium nuclei staining buffer (thaw at RT before, vortex and centrifuge). Incubate 1 min at RT in the dark.
16. Remove all nuclei staining buffer from the well.
17. Add 1,000 μl PBS-T and incubate for 1 min at RT in the dark. Repeat two more times (three in total).
18. Add 1,000 μl PBS-T.

**Table 6.** Drying protocol (for thermal cycler C1000 Touch, Bio-Rad)

| Lid temperature | Reaction volume |  |
| --- | --- | --- |
| 37°C | 100µL |  |
| Step | Temperature | Run time (hh:mm:ss) |
| Pre-equilibrate | 37°C | Hold |
| Drying | 37°C | 00:05:00 |

After this step, the cassettes with slides are ready to be loaded into Xenium Analyzer. Otherwise, slides with the cassette lids can be stored at 4°C in the dark for 1 week. For the best performance it is recommended to replace the PBS-T with a freshly made PBS-T buffer every 3 days, before and after imaging on Xenium Analyzer. Storing slides for more than recommended time risks decrease in assay sensitivity.

#### 3.6 Imaging using Xenium Analyzer

The Xenium Analyzer is initialised (∼10 min) and then all buffers and consumables are loaded (∼5 min) (**Fig. 2C**). Next, the slide scanning is initiated (∼50 min). After scanning is successfully completed, the regions of interest to be imaged are carefully selected manually. The final run time depends on several factors, most critically (i) imaged tissue area, (ii) number of genes in the panel, and (iii) usage of cell segmentation staining. The average Xenium Analyzer run time (imaging and analysis) for our TMAs was 32 hours (software v.3.2.1.2). Selecting each TMA core as a region of interest for imaging (thus reducing the area without tissue) can reduce the run time.

1. Prepare and transfer buffers to the four empty bottles from the Xenium decoding consumables kit.
2. Prepare module A on the day of the run: invert the plate in the intact foil 20x. Centrifuge 2580 rcf for 10 min at RT. If the run is delayed, store it on ice.
3. Prepare module B and cell segmentation module: thaw the plate in original packaging at 4°C for 16–72 h or at 37°C for 2.5 hours in a water bath. Remove plates and equilibrate at RT for 30 min. Invert plates to mix 20x. Centrifuge at 300 rcf for 1 min at RT.
4. On the Xenium Analyzer interface press “Start new run”. Input “Run name”. Tick the “Cell segmentation” option. Add any additional details about the cassettes. Select probe panel. If using a custom panel or an add-on panel, ensure the json format files have been uploaded. To upload a new custom or add-on panel from the USB drive, insert USB into the USB port on the Xenium Analysis computer. Click “Upload” on the instrument touchscreen and select panel “json” file. Confirm the desired panel is selected and click “Continue”.
5. Follow the on screen instructions.
6. Place the module A, module B, and the cell segmentation module into their respective positions.
7. Replace standard bottle caps with Xenium buffer caps (included in the Xenium decoding consumables kit).
8. Place bottles in the bottle carrier in the designated order.
9. Push the bottle carrier caps down to the top of the bottles to seal and load in the appropriate positions.
10. Place the empty waste bottle, pipette tip rack, extraction tip, objective wetting consumable, and waste tip tray into their respective positions.
11. Clean the cassette carrier. Spray 70% isopropanol onto a lint-free laboratory wipe and clean the surface of the carrier.
12. Clean the bottom of the slide surface with 70% isopropanol using a lint-free laboratory wipe without spilling the storage buffer. Confirm the bottom of the slide bottom is clean and dry. This step is crucial for accurate imaging; any dust or debris on the bottom of the slides will impact the performance.
13. Fully open the cassette carrier lid. Place the cassette into the carrier without a lid. Close the cassette carrier lid until it clicks into place.
14. Confirm all consumables are loaded correctly and click “Continue”. Close the instrument front panel.
15. Instrument will begin by sample scanning for 50 min.
16. After the initial scanning, the sample area and related options are shown. Carefully select the regions of interest, which can either be a single ROI area surrounding all tissue cores or multiple ROIs each containing one TMA core. Selecting each TMA core individually, thus limiting the area of empty fields, can improve the run time efficiency.
17. After the run has finished, clean the instrument. Remove consumables and discard solid and liquid waste:

- Xenium cassettes
- waste bottle (discard liquid and rinse distilled water)
- reagent bottles
- reagent plates (module A, module B, and Cell segmentation module)
18. The primary summary quality control (QC) is available to check on the Xenium Analyzer screen. The Xenium Onboard Analysis pipeline, which runs within Xenium Analyser, outputs a summary QC report in HTML format. The report contains summary metrics and automated secondary analysis results. Any warnings are issued at the top of the report (see Section 6 for further details).

### 6. Xenium workflow: data outputs and quality control

After a Xenium run, Xenium Onboard Analysis generates a set of different output files and folders, some of which are described in detail below. The fastest way to view and assess the quality of a Xenium run is to open the summary HTML file, which contains key metrics and spatial information and can be displayed in any web browser.

Additionally, the main output data from tissue level to single cell and transcript resolution, can be visualised within Xenium Explorer software. Processed matrices including cell location, morphology, and segmentation information, as well as transcript location and gene expression, are readily available within the output bundle. Raw data consisting of decoded transcript counts together with the morphology images of the experiment are also provided. In addition to this, and as part of the Xenium Onboard analysis, each Xenium experiment or slide will generate a set of automated downstream analysis files, some of which can be visualised within the summary HTML file and are commented below. Note that output files may change according to the software version used and therefore their contents or format may vary.

1. **Summary HTML file** (*analysis_summary.html*). This file is relevant to assess the quality of the Xenium run at a glance. It contains 5 different sections:

- *Summary*: in addition to Xenium run metadata, this tab contains the total number of cells and high quality transcripts detected, the median number of transcripts per cell, and an interactive heatmap to visualise the density of the transcripts and their quality across the tissue, as well as the negative controls (note this is aggregated data per 20 µm or 80 µm bins) (**Fig. 3A**). In the case of TMAs, where we may observe variability in the RNA integrity between different samples, the heatmap can be a preliminary indication of the performance of the workflow across cores. Considering this variability, and also the fact that these metrics rely on different biological (e.g., tissue type) and technical factors (e.g., panel size), it is challenging to define a threshold for a successful Xenium run. Comparing this data, together with the performance of downstream analyses, across different tissue sections and samples within a given experiment can help to estimate what the boundaries are in a particular scenario. Nevertheless, if any of the metrics fall below 10x defined quality thresholds, warnings and error flags will appear on top of this tab.
- *Decoding*: this tab shows the total number of transcripts from all genes on the Xenium run, including those in the base (pre-designed) and the add-on custom panel, that have been identified or “decoded”, together with their Phred quality scores. Those defined as high quality transcripts have a Phred quality score above 20. The higher the number of high quality transcripts, the higher the quality of the data. The total gene expression per gene, measured as the number of high-quality transcripts for every gene in the pre-designed or the add-on panel, is also shown (Counts per Gene). Marker genes in the tissue should appear at the top of the gene ranking. When assessing the quality and expression level metrics across genes (Gene-Specific Transcript Quality), separation between genuine genes and negative controls will indicate good quality data (**Fig. 3B**). According to 10x guidelines, genes should generally show average quality scores above 20, whereas controls will have scores below 20. Additional information about the negative controls, such as the estimated number of false positive decoded transcripts per cell, is also shown and can be used to evaluate the run in more detail.
- *Cell segmentation*: using this tab, we can determine the number of cells that have been identified using the multimodal cell segmentation algorithm. This uses three different segmentation approaches based on ATP1A1/CD45/E-Cadherin-based (membrane), 18S-based segmentation (RNA), and DAPI (nuclear) staining. The final segmentation result for each cell is prioritised in this order: (i) membrane staining, (ii) expansion from nucleus to the interior RNA stain edge, or (iii) isotropic nuclear expansion of 5 μm. Similar to other QC metrics shown in this tab, including cell size distribution and the number of transcripts detected per cell, the fraction of cells detected by each algorithm will vary according to the tissue type and cellular composition, and therefore these metrics need to be assessed on a tissue- and experiment-specific basis.
- *Analysis*: Similarly to some of the data shown before, this tab will show the spatial distribution of transcript counts, but now across individual cells. As part of Xenium Onboard Analysis, the algorithms will generate graph-based clustering assignments based on gene expression per cell. The spatial distribution of the clusters as well as their UMAP projections of cells is displayed. In the scenario of TMAs, and particularly when samples in the cohort are split between different Xenium slides and runs, this automated analysis will unlikely recapitulate the cell clusters in the whole dataset. Carrying out cell clustering across all Xenium slides is therefore recommended as part of the post-processing steps.
- *Image QC*: this tab shows the fluorescence signals generated during Xenium imaging, as well as those generated by the cell segmentation markers if available. These thumbnail plots show fluorescence intensity across imaging cycles (rows) and channels or colours (columns). Quality data will contain different intensities across cycles and channels, reflecting good signal specificity. If a particular tissue area consistently shows high fluorescence intensity across cycles and channels, it might be a sign of autofluorescence. This could affect transcript detection and impact downstream analysis.
2. **Experiment file** (*experiment.xenium*).This file contains metadata of the Xenium run together with pointers to the actual output data. The experiment file can be loaded into Xenium Explorer, the software provided by 10x (currently available for Windows and Mac users), to interactively visualise the outputs of Xenium experiment. Note that in order for Xenium Explorer to successfully retrieve and display all the data in the run, the experiment file needs to be placed within its original output folder. Once the experiment file has been loaded into Xenium Explorer, users can visualise and interact with tissue morphology images, including DAPI staining for nuclei and cell segmentation markers staining if available (**Fig. 3C-D**), together with segmented cells (**Fig. 3E**) and transcripts signals (**Fig. 3F**). Xenium Explorer allows for extensive customisation to tailor the representation of the information (e.g., overlapping of layers, changes in colours, and signal normalisation) at different scales ranging from whole slide level to intracellular resolution. Note that the Xenium Explorer can import external cell segmentation and cell clustering data. This is particularly convenient for TMAs because the automated clustering generated by Xenium Onboard Analysis will show slide-specific clustering and thus may need to be replaced by a tailored approach using the full set of slides in a project, as discussed above. Xenium Explorer can also be used to export transcript or cell data from manually selected areas within the tissue. Adjacent H&E (or H&E post-Xenium, see Section 8) and immunofluorescence sections can be uploaded into the application, which can help with annotation and further analyses of Xenium slides.

**Figure 3.**
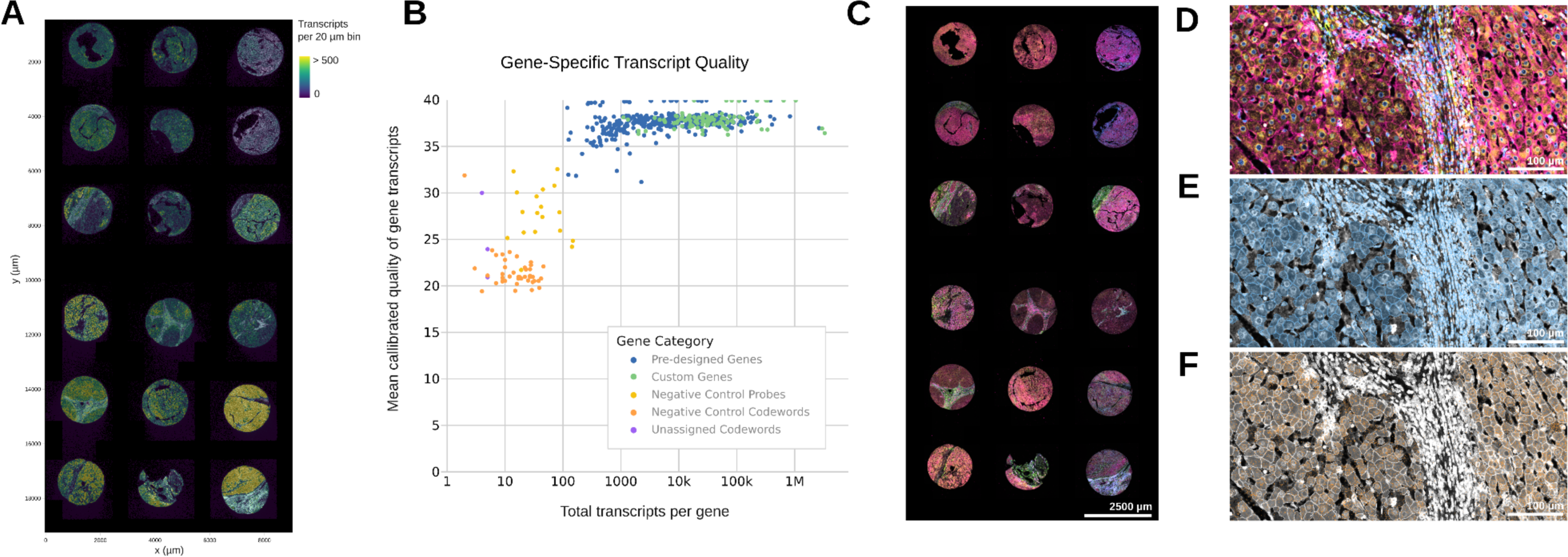
Xenium outputs. **(A)** Heatmap depicting the number of transcripts per 20 μm bin across cores within a Xenium slide. Brighter colours show higher transcript density. The x and y axis show the slide width and height coordinates in μm, respectively. Figure adapted from an output summary HTML file (analysis_summary.html). **(B)** Scatter plot showing transcript quality scores (y axis) versus number of transcripts per gene (x axis) for pre-designed base panel genes, custom add-on genes, and negative controls for one Xenium slide. Figure adapted from the same output summary HTML file (analysis_summary.html) shown in A. **(C)** Visualisation of the Xenium slide shown in A within Xenium Explorer. Morphology images showing DAPI staining in blue (nucleus), ATP1A1, E-cadherin, and CD45 staining in red (membrane), 18S ribosomal RNA staining in yellow (cytoplasm), and alphaSMA and vimentin staining green (cytoskeleton). **(D)** Overlap of morphology images at higher magnification using the same colours as in C. **(E)** Depiction of segmented cell boundaries (blue) for the same tissue area shown in D. Morphology images are overlapped below and shown in greyscale. **(F)** Transferrin (TF) transcripts (coloured orange) are found in hepatocytes. TF transcripts are displayed on top of morphology images (greyscale) and segmented cell boundaries (white) as shown in D and E, respectively.

### 7. Post Xenium data analysis

The data analysis steps after Xenium will be determined by the specific experiment and research question. The first steps will involve quality control assessment; provided that the Xenium workflow results are satisfactory (Section 6), an optional aspect to explore is the performance of cell segmentation outputs. In addition to the automated cell segmentation provided by Xenium Onboard Analysis pipeline, Xenium Ranger (10x) can be used to re-run cell segmentation and incorporate the outputs into the previously generated data. After quality cell segmentation data is obtained, the next steps of Xenium data analysis will be carried out using external methods that are not currently supported by 10x. To date, multiple programmatic tools and libraries in common programming languages have been developed to analyse Xenium data.

Some of these include *Seurat* for R^8^ and Squidpy for Python^9^. In general, Xenium data-analysis pipelines can be divided into three general steps: 1) Data normalisation, filtering, and batch correction at the Xenium slide level; 2) Annotation and data integration. In the case of TMAs, it is key to annotate each tissue core within a Xenium slide with the original tissue metadata (e.g., donor, tissue type, etc). We achieved this by matching post-Xenium H&E and whole TMA adjacent normal H&E tissue sections, where the core location and orientation was maintained. Cells in each core can be segmented and their cell identifiers exported within Xenium Explorer. Once this is achieved, the output data from all Xenium slides in the experiment is integrated and further normalisation is applied, if required; 3) Single cell and spatial data analysis. This part of the analysis will vary according to the specific project. The main steps will include the identification of the cell types and their spatial distribution across multiple scales within the tissue.

### 8. Post Xenium H&E staining and scanning

In order to improve histological comparisons between post-Xenium and adjacent *normal tissues, as well as to* enhance reproducibility of H&E stains, both post-Xenium and *adjacent normal* slides were stained using an autostainer.

1. *Adjacent normal* slides can be stained immediately after baking (Section 4).
2. After a Xenium run, post-Xenium slides could be immediately used for H&E staining. Alternatively, slides can be stored in PBS-T (1000 μl) prior to H&E staining (the PBS-T needs to be refreshed every 3 days).
3. Optional step for post-Xenium slides: quencher removal can be performed prior to H&E staining if the downstream workflow requires it.
  - Immerse in freshly made quencher removal solution for 10 min.
  - 3 x wash in Milli-Q water, 1 min each.
4. H&E staining performed on an autostainer pre-programmed with an FFPE H&E protocol, including coverslipping in DPX.
5. Post-Xenium H&E and adjacent sections H&E stainings are scanned at desired magnification (we used 40X magnification for high resolution).

### 9. Post-Xenium slide and adjacent normal slide storage

Post-Xenium slides can be stored at room temperature in a desiccator, ready for potential future applications, such as immunohistochemistry or immunofluorescence. Note that this is experimental. Please, refer to 10x protocols for future advice on this regard.

### 10. Data storage

Each Xenium run generates approximately 60Gb of data. Specific data storage requirements may be needed according to the local regulations applied to the data generated from each particular experiment (e.g., human data anonymisation).

## Notes

1. Most TMAs have a larger area than the Xenium slide ‘sample area’. Creating a section to fit within this capture area can be achieved in a number of ways. For pre-existing TMAs, a full tissue section can be taken and then the correct region can be cut out from this, or the tissue block can be scored such that the resulting section is the correct size. In this case, we chose the former ‘cutting out’ method to ensure we preserved the integrity of the TMA block for future use. Alternatively a new TMA could be built to exactly match the size of the sample area.
2. Pre-Xenium sectioning optimisation is recommended to achieve high quality sections for the specific tissue type being used.

- Composition of the ice tray for the tissue type, and time for the block to be chilled: Ice that is too dry or blocks that are chilled for too long can cause tissue cracking. Wet ice or under-chilled blocks can lead to holes in the tissue or inconsistent section thickness. For liver TMAs, wet ice for a relatively short time was found to be optimal.
- Water bath temperature: higher temperatures can lead to tissue exploding on the bath, whereas lower temperatures can lead to wrinkled tissue. For liver TMAs ∼45°C was found to be optimal.
- Section thickness: Recommended section thickness for FFPE Xenium is 5 μm, but this can be altered if favourable for the tissue type. TMA cores, especially of liver tissue, have a tendency to roll up during sectioning. This can be exacerbated when section thickness is increased. To test this, the number of rolled up cores were assessed at 4 μm and 5 μm section thickness. The difference was not deemed significant enough to deviate from the protocol recommendation of 5 μm.
3. Passable section quality is subjective and should be agreed up-front, balancing the difficulty of the block being sectioned with the desired outcome. TMAs present a unique challenge with the possibility of individual cores missing or being damaged. Acceptable quality level cannot be prescribed and should take into account the specifics of the TMA such as the presence of duplicated cores, or the number of sections previously taken from the block.
4. The time it takes a section to de-wrinkle and flatten on a water bath is tissue dependent. This can be tested and optimised prior to Xenium sectioning and a lamp can be used to help recognise when the section is uniformly flat.
5. Care should be taken to maintain the orientation of the region as it is being cut out of the rest of the section. If possible, keep the TMA outer orientation cores in place so that if the region is rotated or flipped, it can be re-orientated. If outer orientation cores need to be cut away, extreme care must be taken to keep track of which way round the region needs to be.
6. The 10x Xenium tissue slide preparation including Cell Segmentation step takes three days (**Fig. 2C**). Two slides can be imaged and analysed on the Xenium Analyzer at the same time. It is possible to prepare four tissue slides at the same time and image two slides while keeping the other two slides in the fridge with PBS-T and then load the next two slides straight after the first two slides imaging step is completed.
7. Throughout the Xenium workflow, all reagents should be added and aspirated only when slides are placed in the cassette and on a flat work surface. Avoid contact with tissue, add reagents along the side of the well, avoiding bubbles. The best place to add the reagent is the corner of the well. To remove the reagent, it is more effective to tilt the cassette slightly and use one of the bottom corners. If a bubble forms, carefully puncture with a p10 pipette tip without touching the surface of the slide. Always change tips after each reagent as well as after each slide. The entire surface of the tissue must be covered by the added reagent or buffer, when removing and adding reagents work with one slide at the time to avoid drying out the tissue. Work quickly and ensure reagents are dispensed evenly across tissues during incubation and wash steps throughout the workflow to prevent drying out of tissues. If the reagent is not spread evenly, gently tap the cassette on a flat surface several times.
8. The Xenium thermocycler adaptor should always be in uniform contact with the thermal cycler block surface. Clean the adaptor before use with a lint-free wipe and ethanol. A tight fit between the Xenium cassette and the adaptor, as well as the thermal cycler lid and the cassette lid, is critical during incubation. Make sure you close the lid tightly until you hear an audible click. Tightening the cassette too tightly (after click) may cause the slide to crack. Ensure the thermal cycler has reached an appropriate temperature before starting incubation. There are few critical steps during Xenium workflow where after adding the reagent onto the slides the immediate start of the thermocycler program is critical for an efficient reaction.

## Acknowledgements

We thank the Cancer Research UK Cambridge Institute Core facilities for their valuable contribution: Genomics (Rachel Barnes, Alex Deamer, and Maribel Venezuela for performing Xenium workflows and generating Xenium results); Histopathology and ISH (Sophie Dickinson, Louise Howard, and Julia Jones for technical histology support), and Bioinformatics; and Matt Hoare for assistance with curation of the clinical cohort. This work was supported by grants to S.J.A: Cancer Research UK (Clinician Scientist Fellowship RCCCSF-May23/100001), the Pathological Society of Great Britain and Ireland (3903171 and JSPS CLSG 2020-10-04), the Academy of Medical Sciences (SGL023\1074), and the Medical Research Council (MC_PC_24012).

This is a preprint of the following chapter: C. Arnedo-Pac *et al*, Spatial single cell RNA profiling of human liver FFPE tissue microarrays using Xenium In Situ; an invited submission to In Situ Hybridization Protocols, Sixth Edition, edited by Boye Schnack Nielsen and Julia Jones, Springer Nature. It is the version of the authors’ manuscript prior to acceptance for publication and has not undergone editorial and/or peer review on behalf of the Publisher.

## Notes

### Competing Interest Statement

The authors have declared no competing interest.

